# VC-RDAgent: An efficient rare disease diagnosis agent via virtual case construction informed by hybrid statistical-metric and hyperbolic-semantic prioritization

**DOI:** 10.64898/2026.02.09.702153

**Authors:** Yang Liu, Honglei Li, Peng Jiang, Lizhen Wu, Zhi Xie, Chao Ning, Xiangya Kong, Yayun Wang, Xinlei Zhang, Zechi Huang

**Author notes:** To whom correspondence should be addressed. Email: (Zechi Huang); (Yang Liu).

## Abstract

While Large Language Models (LLMs) have shown promise in clinical decision support, current Retrieval-Augmented Generation (RAG) paradigms face a fundamental bottleneck in rare disease diagnosis: the scarcity, privacy restrictions, and extreme heterogeneity of real-world patient records. This reliance on sparse or inaccessible data leads to a severe “retrieval mismatch,” where the lack of high-quality reference cases causes diagnostic performance to degrade sharply. To break this deadlock, we propose VC-RDAgent, a privacy-preserving and offline-capable framework that decouples diagnostic reasoning from sensitive real-world records by synthesizing virtual standardized cases. The system is powered by VC-Ranker, a multi-dimensional engine that integrates statistical-metric measures with hyperbolic-semantic embeddings to capture deep hierarchical ontology relationships. This approach allows for the dynamic generation of high-fidelity virtual references directly from authoritative knowledge bases. Extensive benchmarking across four diverse datasets demonstrates that VC-RDAgent effectively functions as a “performance equalizer.” It boosts average Top-1 hit rates by 8.7% to 85.9% over zero-case baselines, enabling lightweight open-source models to rival frontier commercial systems. Notably, VC-Ranker alone achieved an aggregate Top-10 hit rate of 0.819, outperforming prior state-of-the-art methods by 6%. By eliminating the dependency on real-time web retrieval and private case sharing, VC-RDAgent provides a scalable, robust, and clinically deployable solution to shorten the diagnostic odyssey, which is made accessible through an intuitive, chat-based web application https://rarellm.service.bio-it.tech/rdagent/.

## 1 Introduction

Rare diseases (RDs) collectively constitute a significant global health challenge, affecting an estimated 350 million individuals worldwide[1]. Although each individual condition exhibits extremely low prevalence, the sheer number of distinct rare diseases, combined with their pronounced phenotypic heterogeneity, creates substantial barriers to accurate and timely diagnosis[2]. These barriers are further exacerbated by extensive phenotypic overlap between rare and common diseases, which often obscures disease-specific signals in routine clinical settings[3]. Furthermore, the multi-system involvement of many rare conditions necessitates a highly coordinated, multidisciplinary approach that is both time-consuming and resource-intensive. As a consequence, patients with rare diseases frequently experience a prolonged “diagnostic odyssey,” characterized by repeated misdiagnoses, delayed interventions, and prolonged uncertainty, ultimately resulting in worsened clinical outcomes and substantial psychological burden[4,5].

To mitigate these challenges, the biomedical community has established a comprehensive ecosystem of structured disease and phenotype knowledge. Authoritative resources such as OMIM[6], Orphanet[7] and Mondo[8], together with the Human Phenotype Ontology (HPO)[9], provide standardized disease definitions, curated phenotype annotations, and interoperable ontological frameworks. These resources have enabled the development of phenotype-driven diagnostic algorithms, including Phen2Disease[10], Phenomizer[11], Phenolyzer[12], LIRICAL-v2.2.1[13] and Exomiser[14], which prioritize candidate diseases by matching patient phenotypes against known disease–phenotype associations. More recent approaches have sought to improve diagnostic performance by integrating additional statistical and machine learning components. Frameworks such as PhenoBrain[15] and PhenoDP[16] combine information-theoretic metrics, graph-based representations, and probabilistic reasoning to refine disease prioritization. In parallel, the emergence of large-scale case-sharing platforms—such as MyGene2 and DECIPHER[17]—has provided access to real-world clinical observations beyond curated databases. These platforms have facilitated case-driven diagnostic tools, including SHEPHERD[18] and PubCaseFinder[19], which better capture phenotypic variability and incomplete presentations commonly observed in clinical practice.

Most recently, large language models (LLMs) have emerged as powerful tools for clinical decision support due to their capacity for semantic reasoning and flexible integration of heterogeneous clinical information[20 – 22]. A widely adopted strategy for applying LLMs to rare disease diagnosis involves Retrieval-Augmented Generation (RAG), specifically utilizing few-shot prompting with retrieved real-world cases, as demonstrated in RareBench[23] and DeepRare[24]. Notably, RareBench reports that such prompting can enable smaller models to approach the performance of frontier models such as GPT-4. However, the effectiveness of real-case–based prompting is fundamentally limited by the nature of real-world clinical data. Available rare disease cases are sparse, unevenly distributed, and highly heterogeneous in quality, reflecting systemic factors such as regional genetic and healthcare disparities, variability in clinician expertise and diagnostic standards, inconsistent phenotype annotation, and privacy constraints that restrict data sharing[25–27]. These limitations make it difficult to obtain standardized, representative reference cases, particularly for ultra-rare or atypical presentations, leading to a frequent retrieval mismatch in which LLM diagnostic performance degrades sharply.

To overcome these limitations, we propose VC-RDAgent, an efficient rare disease diagnosis agent based on systematic virtual case (VC) construction. Instead of relying on sparse and privacy-sensitive real patient records, our framework synthesizes standardized virtual disease cases directly from authoritative biomedical knowledge bases. This strategy addresses the instability of real-case retrieval by establishing a robust ‘knowledge anchor,’ which effectively mitigates the interference of idiosyncratic clinical noise and provides a stable, representative reference space for diagnostic reasoning. At its core, VC-RDAgent is driven by VC-Ranker, a multi-dimensional prioritization engine that integrates information-content specificity, likelihood ratio–based probabilistic evidence, and hyperbolic semantic representations to capture both statistical relevance and hierarchical phenotype–disease relationships. By grounding LLM-based reasoning in these structured, high-fidelity disease archetypes, VC-RDAgent effectively functions as a “performance equalizer”, mitigating retrieval mismatch and ensuring robust diagnostic stability even in data-sparse and highly heterogeneous clinical settings. Furthermore, by decoupling diagnostic inference from private records, our framework offers a scalable, offline-capable and privacy-preserving pathway to shorten the diagnostic odyssey for rare disease patients.

## 2 Methodology

### Overview of VC-RDAgent

The VC-RDAgent is designed as a hierarchical, multi-agent diagnostic system that integrates structured biomedical knowledge with LLM reasoning. The overall architecture is organized into two tightly coupled functional layers: a data orchestration layer and an autonomous diagnostic layer (**Figure 1**).

**Figure 1.**
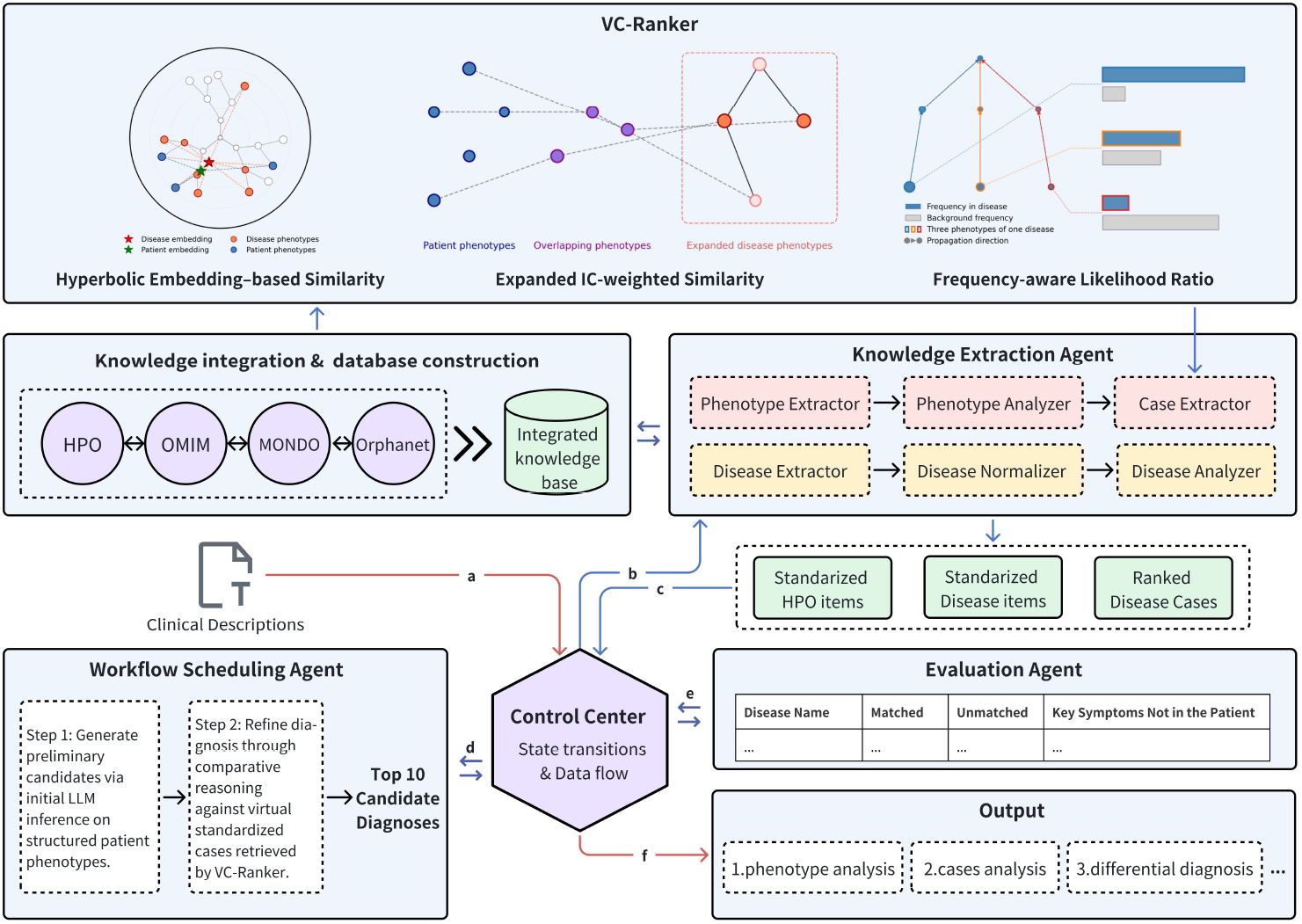
The architecture of VC-RDAgent. a. Raw clinical descriptions are ingested by the control center. b. Input data undergoes phenotype extraction and standardization. c. Structured metadata and virtual cases prioritized by VC-Ranker are retrieved. d. The agent executes the two-stage diagnostic reasoning. e. Diagnostic candidates are evaluated for clinical consistency. f. A comprehensive diagnostic report is generated.

**Figure 2.**
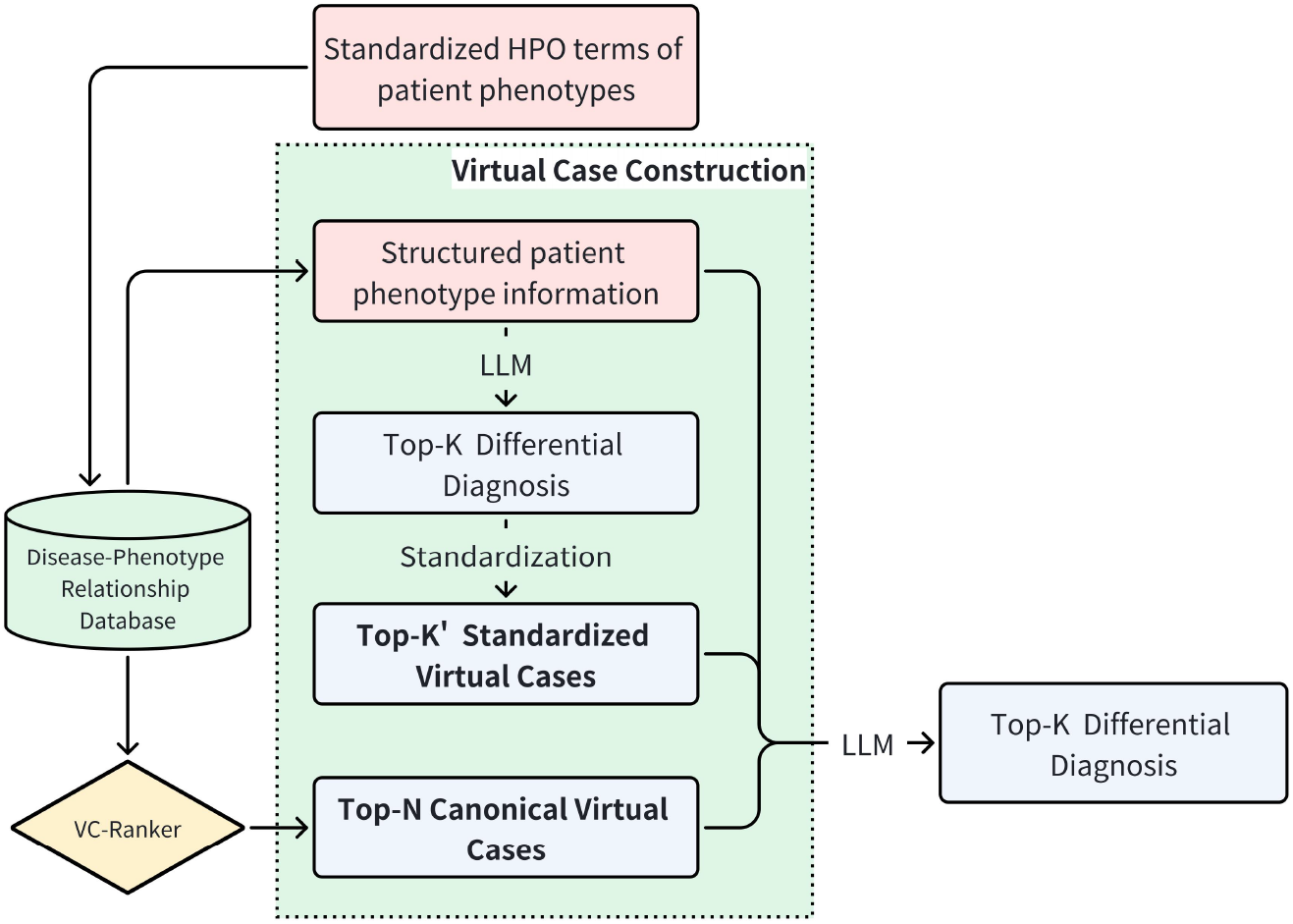
The fundamental architecture of virtual case construction.

The data orchestration layer is responsible for standardizing clinical inputs and constructing a unified disease–phenotype knowledge representation. Raw clinical descriptions are first mapped to standardized HPO terms using FastHPOCR[28] and LLM. In parallel, a knowledge extraction agent integrates curated evidence from HPO, OMIM, Orphanet, and MONDO to construct a structured, cross-ontology disease–phenotype graph.

The autonomous diagnostic layer is governed by a control-center agent responsible for coordinating reasoning states and managing information flow. To support efficient candidate retrieval and phenotype-driven case construction, the system incorporates VC-Ranker, a multi-dimensional disease prioritization engine bridging knowledge orchestration and autonomous reasoning. VC-Ranker introduces a hyperbolic embedding–based semantic similarity module to model latent hierarchical relationships between patient phenotypes and diseases, together with expanded IC-weighted phenotype overlap and frequency-aware likelihood ratios for joint disease prioritization. The resulting ranked disease list supports downstream virtual case construction and comparative diagnostic reasoning within the autonomous diagnostic layer. Within this layer, the workflow scheduling agent executes a structured chain-of-thought (CoT) diagnostic process that jointly considers the standardized patient phenotype profile and a dynamically synthesized set of virtual cases. An independent evaluation agent subsequently performs consistency checks on the generated differential diagnoses, assessing clinical plausibility and suppressing spurious inferences, thereby reducing hallucinations and enhancing diagnostic safety.

### Knowledge integration and database construction

To support automated virtual case construction and multi-agent diagnostic reasoning, we constructed a high-coverage, concept-centric disease–phenotype knowledge base by systematically integrating data from HPO, Orphanet, OMIM, and MONDO. The integration process follows a MONDO-centered alignment strategy, leveraging curated cross-ontology mappings and synonym sets to unify heterogeneous disease identifiers into a single semantic framework.

Each integrated disease entity inherits complementary attributes from the contributing resources. Specifically, HPO provides the standardized phenotype vocabulary and hierarchical structure; OMIM contributes genotypic and pathogenic mechanism annotations; Orphanet supplies expert-curated clinical classifications, genotypic annotations, and rarity tiers; and MONDO ensures semantic consistency and cross-database interoperability. Through this aggregation, each disease entity is associated with a rich, multi-dimensional phenotypic signature that includes both qualitative descriptions and quantitative frequency annotations. This unified architecture enforces consistency across nomenclature, coding systems, and hierarchical relations, enabling direct comparative analysis within a single computational environment. By consolidating the strengths of individual databases while mitigating their respective coverage limitations, the resulting repository offers a more comprehensive and fine-grained representation of the rare disease landscape than any standalone source.

The finalized knowledge base comprises 16,629 disease entities, including 12,208 rare diseases, and contains 252,464 disease–phenotype associations enriched with frequency information. Beyond its role in virtual case construction, this scalable resource supports longitudinal updates and provides a stable foundation for phenotype-driven inference, phenotype–genotype correlation analysis, and future diagnostic system extensions.

### VC-Ranker

Building on the unified knowledge base, we developed VC-Ranker, a multi-dimensional disease prioritization engine that explicitly integrates statistical-metric phenotype similarity and hyperbolic-semantic proximity for candidate disease retrieval in virtual case construction and downstream diagnostic reasoning. Specifically, VC-Ranker combines two complementary statistical paradigms, Expanded IC-weighted phenotype similarity and frequency-aware likelihood ratio, to quantify explicit phenotypic concordance and diagnostic evidence strength, while a hyperbolic embedding based semantic similarity module captures latent hierarchical and semantic relationships beyond direct ontology matching.

#### Expanded IC-weighted phenotype similarity

The first component emphasizes diagnostic specificity by weighting phenotypes according to their information content (IC). The IC of a phenotype *i* is inversely proportional to its prevalence across the integrated disease knowledge base, calculated as:

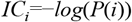

where *P*(*i*) denotes the proportion of diseases annotated with phenotype *i* in the knowledge base. A higher IC value indicates a more specific symptom with greater diagnostic significance. The IC-based similarity score, *S*_*ic*_, is defined as the ratio of the summed IC of overlapping phenotypes to the total IC of the patient’s profile:

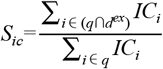

where *q* denotes the patient’s phenotype set, and *d*^*ex*^ represents an expanded set of disease phenotypes including ancestral and unique descendant terms to enhance matching robustness. This metric ensures that rare, highly informative phenotypes contribute proportionally more to the overall similarity score, and that the computation leverages the breadth of the integrated database to improve robustness.

#### Frequency-aware likelihood ratio

The third component of VC-Ranker integrates clinical frequency data through a probabilistic ranking algorithm grounded in the Likelihood Ratio (LR) framework and Bayesian inference. This approach quantifies the diagnostic relevance of each observed phenotype by evaluating its prevalence within a specific candidate disease relative to its background frequency across the entire disease spectrum. For each observed phenotype *i* in a patient, the algorithm calculates an LR score as follows:

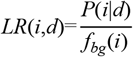

where *P*(*i*|*d*) represents the expected frequency of phenotype *i* in disease *d*, and *f*_*bg*_(*i*) denotes its background frequency. The background frequency is the average over all diseases in *D* of the maximum annotated frequency of phenotype *i* or any of its descendants in each disease *d*, defined as:

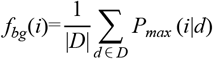

To maximize the utility of the often sparse frequency annotations in public databases, we implement a distance-decay propagation mechanism. This mechanism allows the engine to handle cases where the frequency of a node *i* is not directly recorded for a disease *d* by estimating an induced frequency, *P*_*induced*_(*i*|*d*), derived from its annotated descendants:

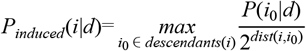

In this equation, *descendants*(*i*) refers to all child nodes of phenotype *i, P*(*i*_0_|*d*) is the directly annotated frequency of a descendant node *i*_0_ in disease *d*, and *dist*(*i,i*_0_) represents the hierarchical distance between phenotype *i* and its descendant *i*_0_. By multiplying the LR values of all independent phenotypic evidence, the system generates a composite likelihood ratio, which is subsequently combined with the prior probability of each disease to calculate a final posterior probability. This probabilistic framework ensures that the ranking is not only driven by the presence of a symptom but also by the quantitative strength of its association with the disease, providing a highly interpretable and evidence-based candidate list.

#### Hyperbolic embedding–based semantic similarity

To capture deep hierarchical and semantic relationships, VC-Ranker incorporates a newly introduced hyperbolic embedding[29] module. This approach leverages hyperbolic space to mirror the exponential branching structure of biological ontologies, allowing for the representation of complex hierarchies in lower dimensions with minimal distortion. The embedding space is constructed from a heterogeneous graph integrating phenotype-disease, phenotype-gene, gene-disease, and phenotype-phenotype associations.

The hyperbolic embeddings are generated by training a poincaré ball model[30], in which each node of the knowledge graph (disease, phenotype, or gene) is represented as a point constrained within the unit hyperbolic ball. Edge types, including hierarchical relationships, causal links, and disease–phenotype associations, are assigned weights reflecting information content and observed frequencies, which guide the contrastive learning objective. The training procedure employs Riemannian optimization[31] to respect the manifold geometry, and negative sampling is used to enforce separation of unrelated nodes. For aggregating multiple phenotype embeddings, an Einstein midpoint operation in the poincaré ball is used to compute a weighted centroid, allowing patient phenotypes to be represented as a single hyperbolic vector while preserving hierarchical distances.

Rather than using simple vector averaging, we implement an asymmetric “best-match” strategy to align patient symptoms with disease signatures. Let *E*_*i*_ be the embedding vector for phenotype *i* and *IC*_*i*_ be its information content. For a patient phenotype set *q* and a candidate disease phenotype set *d*, the semantic similarity score *S*_*emb*_ is defined as:

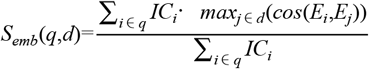

where *cos*(*E*_*i*_,*E*_*j*_) denotes the cosine similarity between the embeddings. This formulation ensures that for each clinical manifestation in the patient, the engine identifies the most relevant corresponding feature in the disease profile, effectively mitigating noise from unrelated phenotypes while emphasizing critical semantic matches.

Finally, the outputs from the IC-based similarity, phenotypic likelihood ratio and hyperbolic semantic embedding modules are aggregated using a rank-based consensus method to produce the definitive prioritized diagnostic results.

### Virtual case construction

Virtual case construction serves as the cognitive core of the workflow scheduling agent, orchestrating the transformation of structured clinical data into a robust, knowledge-grounded environment for diagnostic reasoning. By synthesizing high-density “Virtual Standardized Cases” directly from the integrated knowledge base, the framework decouples diagnostic reasoning from sparse and privacy-sensitive real-world records. These virtual cases are dynamically constructed from two distinct sources: prioritization via VC-Ranker and standardized heuristic candidates generated by the LLM.

The first source of virtual cases is generated by performing a global similarity search across the entire knowledge base using the VC-Ranker engine. The engine identifies the Top-N (N=50) diseases that best align with the patient’s phenotype profile, leveraging its integrated statistical and semantic prioritization modules. These prioritized candidates are transformed into virtual standardized cases, where each entry represents an expert-curated, population-level disease archetype. By providing explicit phenotypic frequencies and structural metadata for each retrieved candidate, this process establishes a stable and representative reference space for comparative reasoning.

In parallel, the framework incorporates a heuristic pathway to capture the LLM’s internal medical knowledge. Upon receiving the standardized patient phenotypes, the workflow scheduling agent performs an initial diagnostic inference to yield a preliminary Top-K (K=10) differential diagnosis list. To ensure clinical rigor and prevent hallucinations, these raw model outputs undergo a semantic normalization process, where candidates are mapped back to the standardized disease entities (e.g., MONDO, Orphanet, or OMIM identifiers) in the integrated database. Only those diseases successfully resolved to a database identifier are retained, resulting in a refined set of Top-K′ (K’ ≤ 10) candidates. These diseases are then enriched with their corresponding structured phenotypic signatures, effectively converting the model’s heuristic hypotheses into standardized virtual cases.

The final augmented prompt integrates the patient’s clinical metadata with these retrieved and standardized virtual cases. This structured evidence space anchors the agent’s reasoning in authoritative biomedical knowledge, mitigating the impact of phenotypic heterogeneity and data sparsity. Consequently, the final diagnostic output emerges from comparative reasoning over synthesized disease archetypes rather than fragile dependence on incomplete real-world case retrieval.

### Benchmark dataset

The VC-RDAgent was systematically evaluated using four real-world rare disease repositories with diverse clinical profiles and data origins. PUMCH-ADM and HMS represent high-quality institutional datasets, comprising 75 cases (16 diseases) from Peking Union Medical College Hospital and 88 cases (39 diseases) from Hannover Medical School, respectively[15]. As single-center records, these datasets exhibit high internal consistency in diagnostic protocols and phenotype mapping but may reflect specific regional genetic backgrounds and clinical practices.

To assess the model’s robustness against data heterogeneity, we included the MyGene2[18] and LIRICAL[15] datasets. MyGene2 (146 cases, 55 diseases) consists of patient-driven data from a global sharing platform managed by the University of Washington. Unlike institutional records, this repository introduces significant phenotypic noise and variability characteristic of family-reported health information, testing the framework’s resilience to non-expert descriptions. In contrast, the LIRICAL dataset, curated by the Jackson Laboratory for Genomic Medicine, serves as a literature-based gold standard with 370 cases covering 252 diseases. These cases, extracted from peer-reviewed clinical reports, provide highly structured and prototypical disease representations. By combining institutional, patient-reported, and literature-derived cases, this benchmark suite ensures a comprehensive assessment of VC-RDAgent’s generalizability and stability across varying levels of data quality and clinical complexity.

## 3 Results and Discussion

### Performance of the VC-Ranker engine

We first evaluated the Top-K (K=1, 5, 10) hit rates of VC-Ranker across four shared real-world rare disease case repositories, comparing its performance with representative phenotype-driven prioritization methods including LIRICAL-v2.2.1, Exomiser, and PhenoDP. While LIRICAL-v2.2.1 and Exomiser utilize traditional statistical and variant-annotation frameworks, PhenoDP represents a more recent comprehensive multi-strategy approach.

As illustrated in the experimental results, VC-Ranker achieved performance that was either comparable to or exceeded the current state-of-the-art methods across the PUMCH-ADM, HMS, and LIRICAL datasets. It is worth noting that all methods tend to perform particularly well on the LIRICAL dataset, which is curated from peer-reviewed literature and provides highly structured, prototypical disease cases. In contrast, in the HMS repository, VC-Ranker demonstrated a significant advantage across all metrics, with its Top-1 hit rate improving by over 200% compared to the next best method, PhenoDP. In the PUMCH-ADM dataset, VC-Ranker maintained comparable Top-1 accuracy while consistently outperforming all other methods in Top-5 and Top-10 hit rates. This performance pattern highlights robustness of the hybrid prioritization strategy: by leveraging a deep semantic understanding of phenotype-disease associations, VC-Ranker stabilizes correct diagnoses within top-tier candidates even when atypical features preclude a Top-1 hit. This high fault tolerance provides a reliable diagnostic safety net, ensuring target diseases remain within the clinician’s primary focus.

In the MyGene2 repository, which integrates cases submitted by both patients and researchers and is characterized by non-standardized descriptions and high linguistic noise, LIRICAL-v2.2.1 maintained an advantage in Top-1 accuracy, possibly because its highly optimized likelihood-ratio architecture is exceptionally sensitive to individual high-specificity features. However, the performance gap between VC-Ranker and LIRICAL-v2.2.1 narrowed significantly at the Top-10 level. This further validates the aforementioned robustness: despite the noise or ambiguity in symptom descriptions, VC-Ranker effectively mitigates interference by capturing broad phenotypic associations, successfully recalling the correct diagnosis into the differential list. Collectively, VC-Ranker demonstrates not only high precision on curated clinical data but also exceptional generalization and clinical safety across heterogeneous and noisy data sources.

Summary statistics across the total 679 cases further confirm the comprehensive performance advantages of VC-Ranker. The engine achieved the highest scores across all evaluation metrics, with aggregate Top-1, Top-5, and Top-10 hit rates of 0.546, 0.736, and 0.819, respectively. Notably, its Top-10 performance represents a 6% lead over both PhenoDP and LIRICAL-v2.2.1, and a substantial improvement over Exomiser (0.652). These results establish VC-Ranker as a robust and versatile tool for rare disease prioritization, providing a stable foundation for the subsequent virtual case-enhanced diagnostic reasoning of the LLM across diverse real-world clinical scenarios.

### Structured virtual cases substantially enhance LLM diagnostic accuracy

To quantify the contribution of virtual cases to the diagnostic performance of LLMs, we benchmarked seven models across four diverse clinical datasets (**Table 1**). As illustrated in **Figure 4**, VC-enhanced LLMs (VC-LLMs) demonstrated a profound improvement in diagnostic accuracy compared to the zero-case baseline. Specifically, the average Top-1 hit rate for VC-LLMs increased by approximately 8.7% in PUMCH-ADM, 14.1% in HMS, 79.8% in MyGene2, and 85.9% in LIRICAL. Furthermore, average Top-5 accuracy across the four datasets rose from a zero-case baseline range of 49.2%–69.5% to a VC-enhanced range of 65.7%–81.2%. These results indicate that the virtual case strategy substantially enhances the differential diagnosis capabilities of LLMs, providing more reliable guidance for clinicians, particularly when the model’s internal representation of specific rare diseases is sparse or when critical diagnostic features are underrepresented in the training data.

**Table 1.** Seven LLMs evaluated as rare disease specialists.

| Model | Source | Parameter | Open Source | Mode |
| --- | --- | --- | --- | --- |
| Gemini-2.0-Flash | Google | / | No | thinking |
| GPT-4o-Mini | OpenAI | / | No | no-thinking |
| GPT-oss-120B | OpenAI | 117B | Yes | thinking |
| DeepSeek-Chat-v3.2 | DeepSeek | 685B | Yes | no-thinking |
| Qwen3-8B | Qwen | 8.2B | Yes | thinking |
| MiniMax-M2 | MiniMax | 230B | Yes | thinking |
| GLM4.7 | Zhipu AI | 358B | Yes | thinking |

The integration of LLM reasoning with structured virtual cases also demonstrated a clear advantage over traditional algorithmic approaches. Compared to the standalone VC-Ranker, VC-LLMs achieved higher Top-1 hit rates in the PUMCH-ADM, HMS, and MyGene2 datasets, with improvements of 67.7%, 91.2%, and 4.7%, respectively. This superiority underscores the value of the logical reasoning inherent in LLMs, which can effectively refine diagnostic priorities in scenarios where real-world phenotypic patterns deviate from the idealized descriptions found in standard databases. However, a nuanced exception was observed in the LIRICAL dataset, where VC-LLMs showed a slight decrease in performance compared to VC-Ranker (with an average Top-1 recall of 0.632 versus 0.676, respectively). Given that LIRICAL cases are typically derived from published clinical reports and with highly structured and prototypical cases, our similarity-based algorithms can achieve high precision through direct pattern matching (**Figure 3**). In such instances, the introduction of complex reasoning through an LLM may lead to minor interference from non-critical phenotypic noise, particularly in lower-parameter models. Nevertheless, this does not diminish the overall utility of the virtual cases. Since phenotypic heterogeneity and atypical presentations are common in real-world clinical practice and are challenging for traditional similarity-based methods, as discussed above, the ability of VC-LLMs to maintain robust performance across diverse and incomplete clinical records makes it a more versatile tool for real-world clinical application.

**Figure 3.**
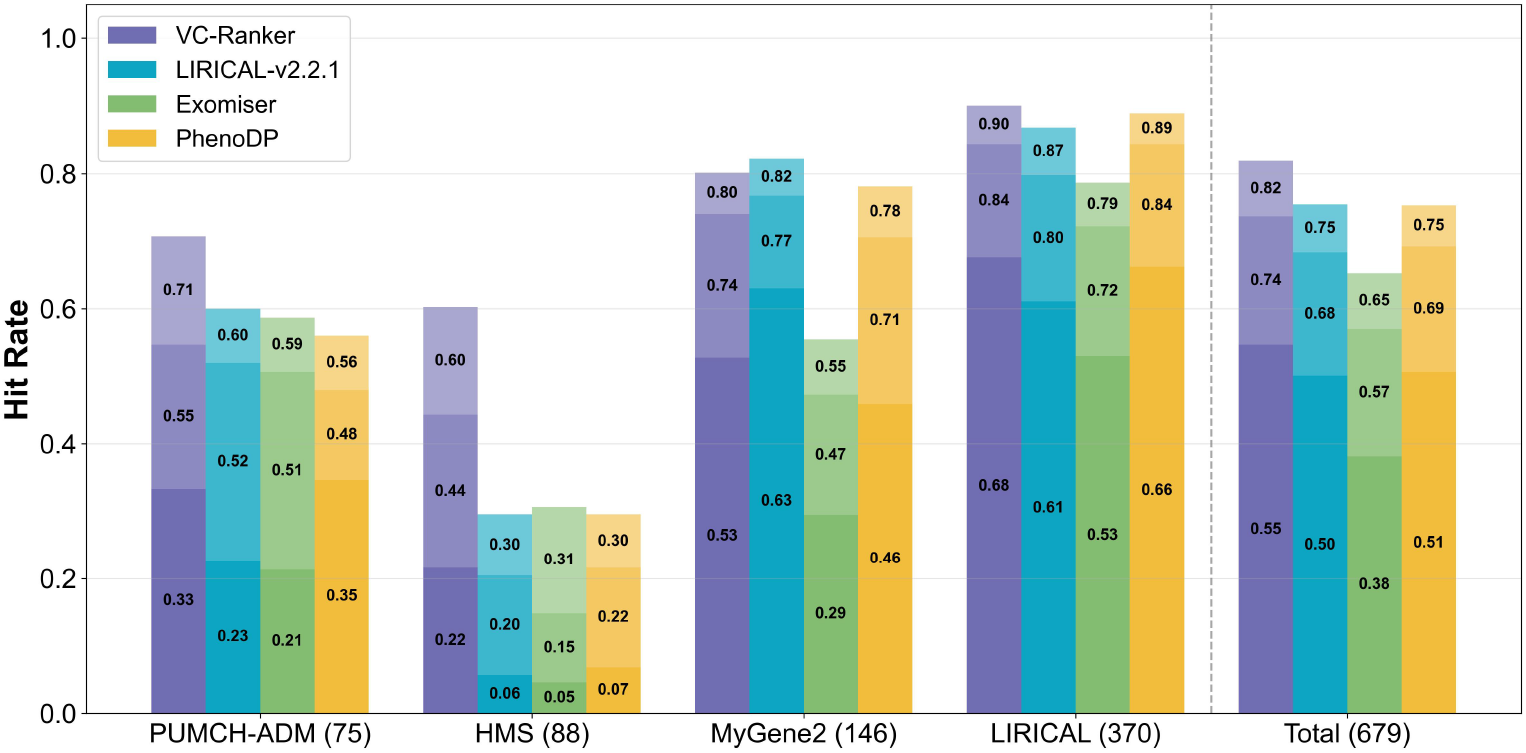
Top-K disease hit rates of VC-Ranker across four real-world rare disease case repositories. Each bar is segmented from dark to light indicating Top-1, Top-5, and Top-10 hit rates.

**Figure 4.**
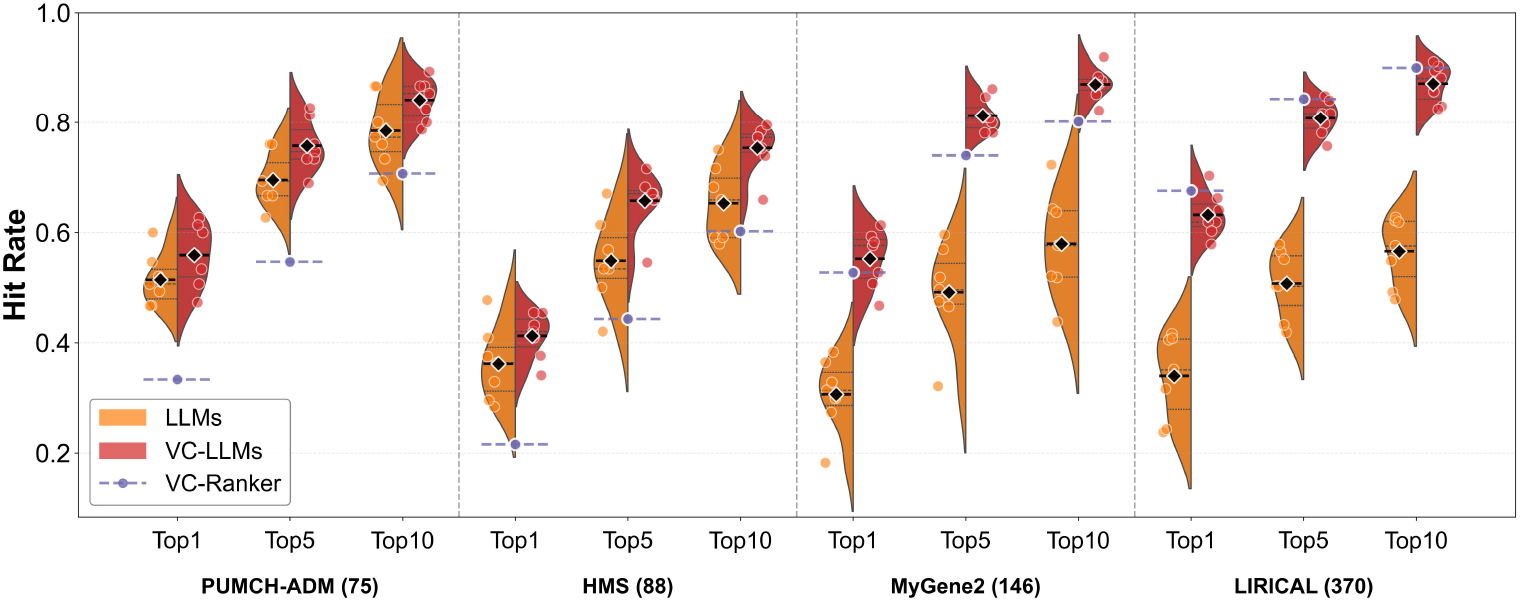
Split violin plots comparing the performance of LLMs and VC-LLMs across four datasets. Individual data points from multiple models are shown as scatter points. Quartiles are indicated by the inner structure of the violins, and mean values are marked with horizontal lines and diamond markers.

### Structured virtual cases harmonizes diagnostic performance across model scales and narrows the frontier gap

Crucially, the virtual cases act as a “performance equalizer,” harmonizing diagnostic efficacy across diverse model architectures. As illustrated in **Figure 5**, the VC-strategy effectively bridged the parameter gap: the 8.2B-parameter Qwen3-8B surpassed the 685B-parameter DeepSeek-v3.2 in PUMCH-ADM and MyGene2, while remaining highly competitive in HMS and LIRICAL within a narrow 3–5% margin of larger peers. Furthermore, the frontier open-source models consistently matched or led proprietary systems across most cohorts: VC-enhanced GLM4.7 and MiniMax-M2 outperformed Gemini-2.0-flash and GPT-4o-mini in PUMCH-ADM and HMS, and remained among the top three performers in MyGene2 and LIRICAL, where Gemini-2.0-Flash maintained a slim lead. This indicates that virtual cases provide a standardized knowledge anchor that allows open-source models to overcome the scale advantages of closed-source commercial models.

**Figure 5.**
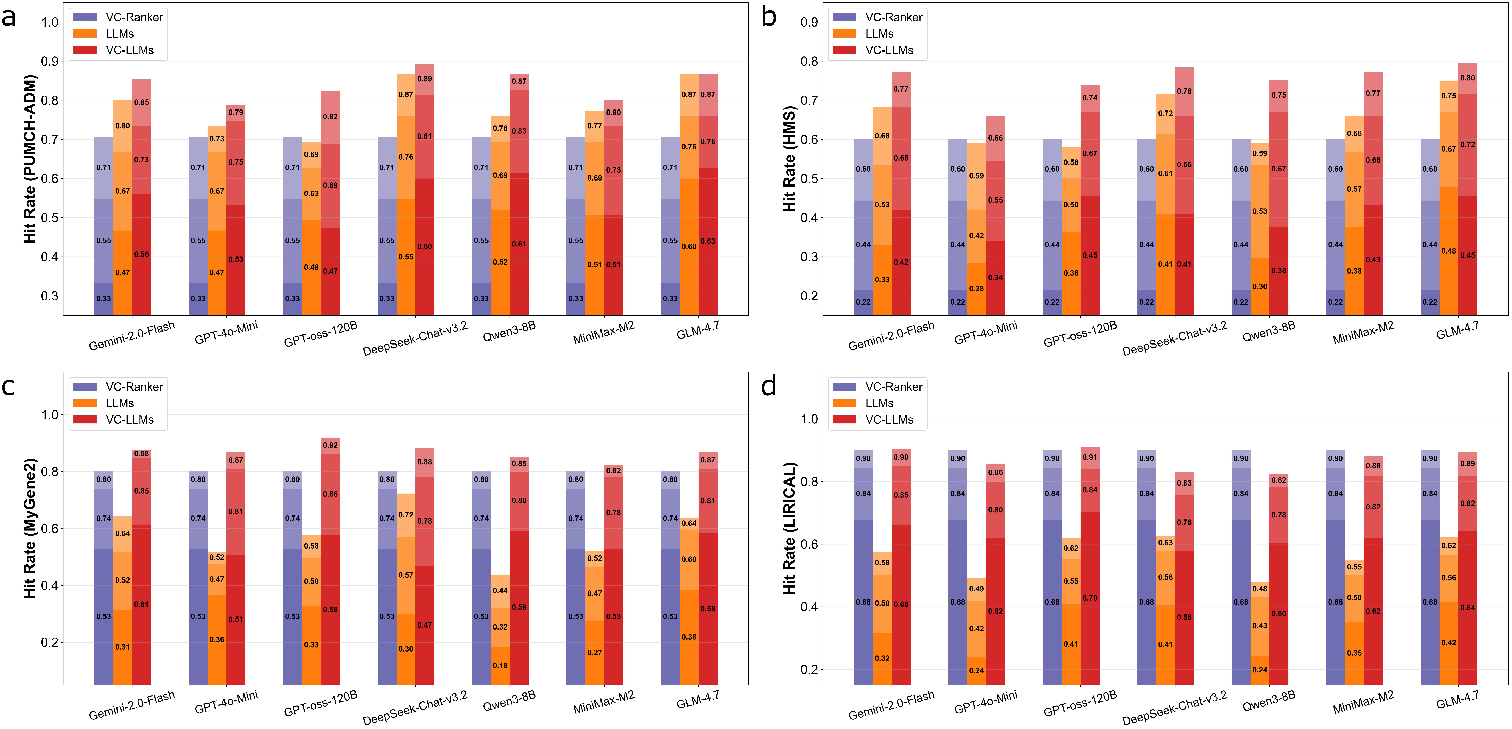
Diagnostic performance of different LLMs on rare disease case libraries.

Beyond bridging performance differences across model architectures, the VC-strategy markedly enhances the stability and consistency of diagnostic performance across heterogeneous clinical settings. By comparing the fluctuations in Top-K hit rates, we quantified stability using the coefficient of variation (CV), defined as *CV*= *σ/μ*, where *σ* and *μ* are the standard deviation and mean of Top-1 hit rates across datasets, respectively. VC-LLMs (**Figure 5**) exhibited the highest degree of robustness against dataset variability: for instance, while the Top-1 hit rate of VC-Ranker (**Figure 3** and **Figure 5**) ranged from 0.216 to 0.676 (CV = 0.47) and zero-case LLMs ranged from 0.18 to 0.6 (CV = 0.26), VC-LLMs maintained a much narrower and more stable range of 0.34 to 0.70 (CV = 0.17). This suggests that the standardized virtual cases serve as a critical “knowledge anchor,” mitigating the impact of varying data quality, regional diagnostic standards, and phenotypic completeness. Consequently, the VC-strategy not only elevates the average diagnostic ceiling but also ensures that the system remains reliable when faced with highly heterogeneous clinical data, providing a scalable and generalizable solution for the “diagnostic odyssey” faced by rare disease patients.

To determine the performance ceiling of the VC-strategy, we compared the average performance of the seven mid-size models (7-LLMs) against the frontier commercial models (GPT-5 and Gemini-3-Pro) on the HMS dataset (**Table 2**). The results demonstrate that the structured virtual cases consistently bolsters diagnostic recall across all model tiers. For both frontier models, VC-LLMs led to a steady expansion of the diagnostic horizon, with Top-10 hit rates reaching 0.830 for GPT-5 and 0.851 for Gemini-3-Pro. While Top-1 gains varied between these top-tier models, the universal improvement in Top-5 and Top-10 metrics indicates that the VC-strategy effectively stabilizes the broader differential diagnosis list, ensuring the correct disease is captured within the primary clinical focus. For the 7-LLMs, the structured virtual cases acted as a systemic performance elevator, significantly narrowing the gap with frontier models. As shown in **Table 2**, the application of virtual cases increased the average Top-10 hit rate for mid-size models from 0.653 to 0.754, effectively approaching the zero-case baseline of GPT-5 and Gemini-3-Pro. Notably, high-parameter open-source models such as deepseek-chat-v3.2 and glm4.7 achieved Top-10 recall of 0.784 and 0.795 respectively with virtual cases, placing them within a narrow 2–4% margin of the zero-case GPT-5 performance and directly comparable to frontier commercial systems.

**Table 2.**
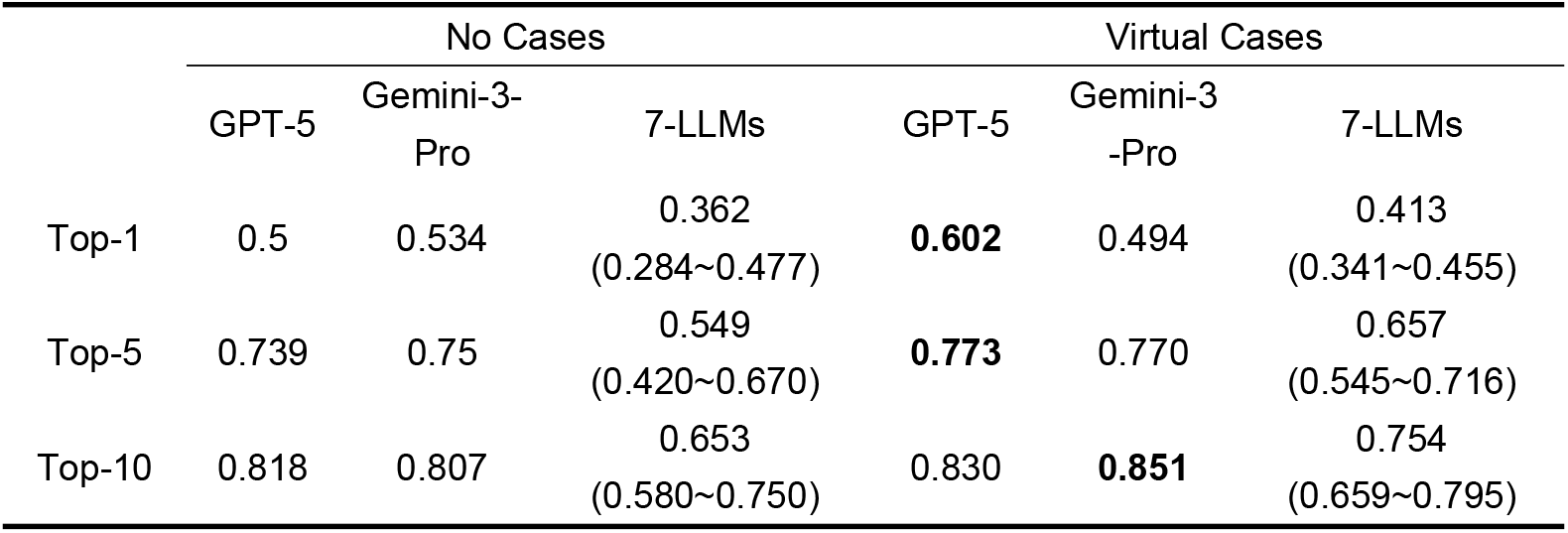
Top-K diagnostic accuracy of GPT-5, Gemini-3-pro and an average of seven LLMs (7-LLMs) under no-cases and VC-strategy in the HMS benchmark.

|  | No Cases |  |  | Virtual Cases |  |  |
| --- | --- | --- | --- | --- | --- | --- |
|  | GPT-5 | Gemini-3-Pro | 7-LLMs | GPT-5 | Gemini-3-Pro | 7-LLMs |
| Top-1 | 0.5 | 0.534 | 0.362<br>(0.284~0.477) | **0.602** | 0.494 | 0.413<br>(0.341~0.455) |
| Top-5 | 0.739 | 0.75 | 0.549<br>(0.420~0.670) | **0.773** | 0.770 | 0.657<br>(0.545~0.716) |
| Top-10 | 0.818 | 0.807 | 0.653<br>(0.580~0.750) | 0.830 | **0.851** | 0.754<br>(0.659~0.795) |

### Competitive diagnostic performance with superior privacy and robustness

A core strength of the VC-strategy lies in its use of “virtual cases” derived from public knowledge bases, rather than relying on sensitive real-patient records or real-time web-crawling. To evaluate whether this privacy-preserving and offline-capable strategy can compete with retrieval-based and search-enhanced systems, we conducted a controlled comparison utilizing Gemini-2.0-Flash, the shared base model evaluated by both methodologies, to ensure a direct and fair assessment (**Figure 6**).

**Figure 6.**
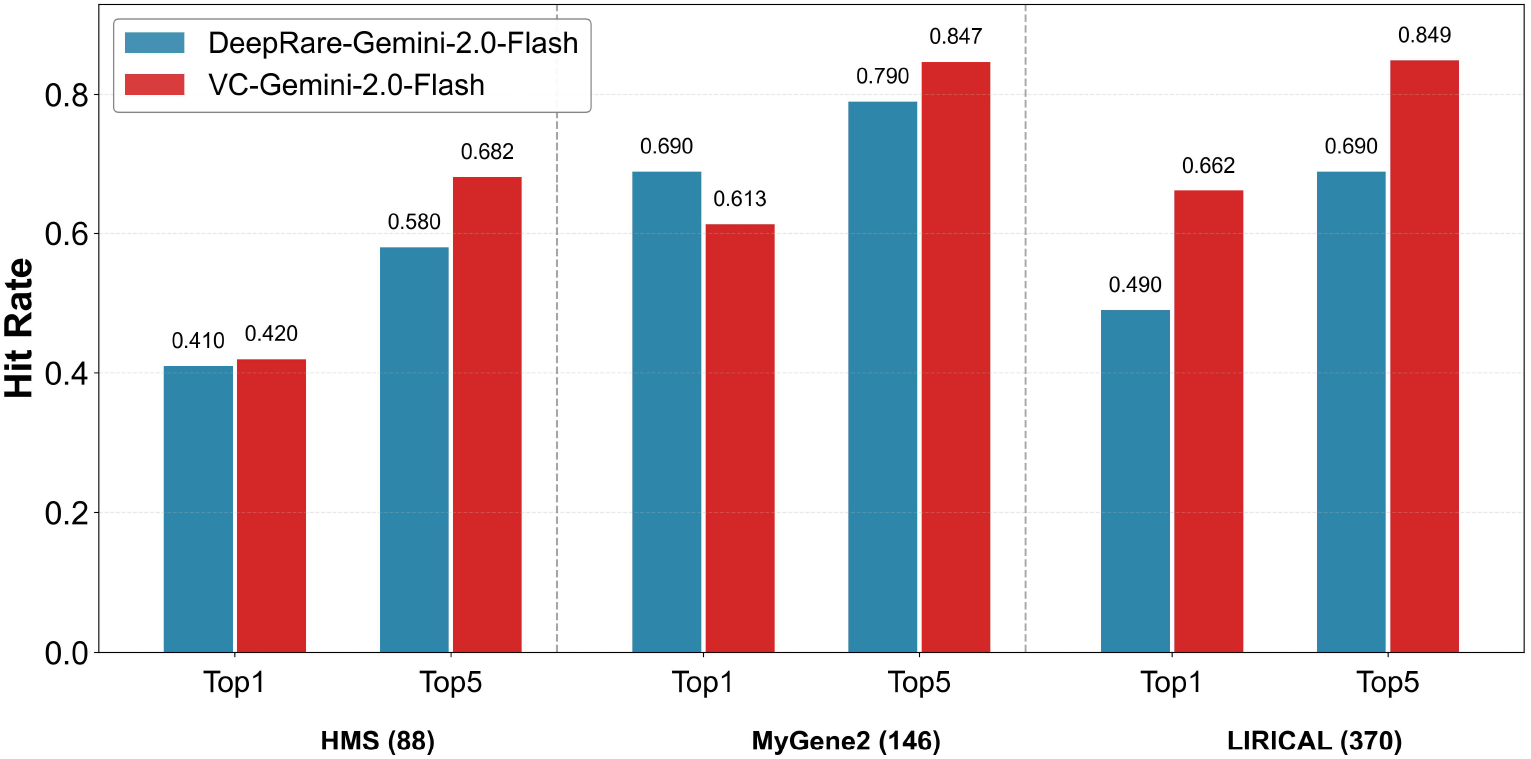
Comparative diagnostic performance of DeepRare-Gemini-2.0-Flash and VC-Gemini-2.0-Flash across three shared rare disease benchmarks. Metrics of DeepRare are cited from its publication[24], while the hit rate of VC-Gemini-2.0-Flash was calculated using identical evaluation protocols to ensure comparability.

The results demonstrate that the synthesized “knowledge anchor” offers superior robustness compared to real-time retrieval. In expert-curated environments (LIRICAL and HMS), VC-Gemini-2.0-Flash consistently outperformed DeepRare, most notably achieving a dominant Top-1 accuracy of 66.2% in LIRICAL and establishing a substantial 10% lead in HMS Top-5 accuracy alongside a marginal Top-1 advantage. This suggests that in clinical contexts with standardized descriptions, representative virtual cases effectively filter out the idiosyncratic noise and confounders often introduced by web-crawled records, allowing the model to converge more accurately. In the MyGene2 dataset, characterized by noisy patient-reported descriptions, DeepRare retained a lead in Top-1 accuracy, likely utilizing web search to match specific idiosyncratic phrasings. Crucially, however, VC-Gemini-2.0-Flash reversed this trend in the broader differential, surpassing DeepRare in Top-5 accuracy by 5.7%. This indicates that while real-time retrieval may occasionally achieve a “first-hit” via keyword matching, the VC-strategy provides a more reliable diagnostic safety net, delivering superior clinical value in a fully offline, privacy-preserving framework.

To further explore the differences between real-world clinical evidence and synthesized virtual knowledge in enhancing model performance, we implemented the real case augmented (RC) strategy. This strategy follows the core dynamic few-shot mechanism proposed in RareBench[23], where the three most similar real cases are retrieved for each test sample to serve as contextual references. However, our implementation differs in two key aspects: the hybrid phenotype-similarity prioritization method (RC-Ranker) and the specific case construction technique used to integrate the retrieved cases into the LLM reasoning. For the comparison, we utilized the lightweight Qwen3-8B model. The results, as shown in **Figure 7**, indicate that the efficacy of RC-strategy versus VC-strategy is highly dependent on the intrinsic characteristics of the case repository.

**Figure 7.**
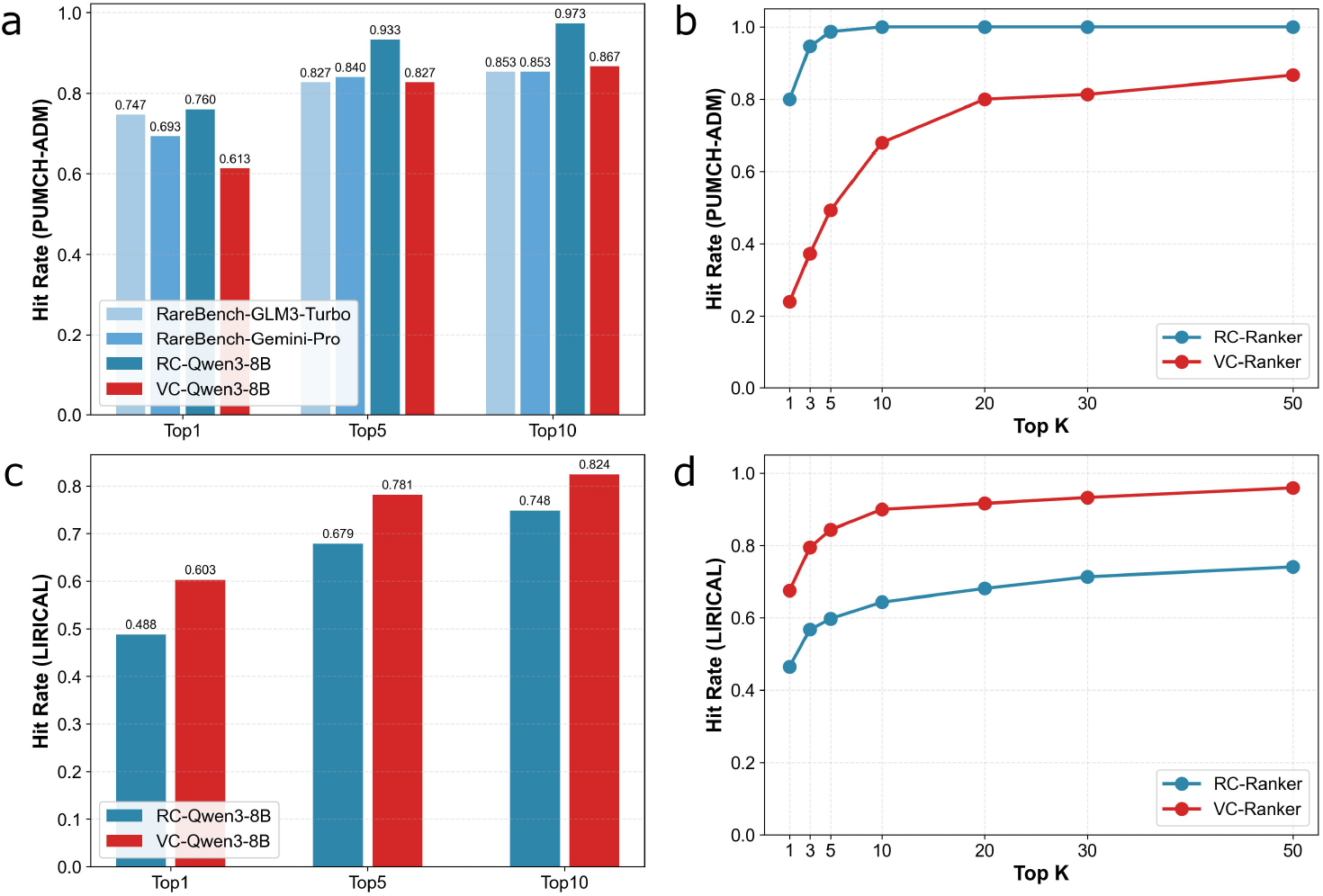
Comparative efficacy of virtual and real case strategies in disease retrieval and LLM-assisted diagnosis. a. Comparison of hit rates between RareBench, RC-Qwen3-8B and VC-Qwen3-8B in PUMCH-ADM dataset. b. Hit rates of VC-Ranker and RC-Ranker across different Top K values in PUMCH-ADM dataset. c. Comparison of hit rates between RC-Qwen3-8B and VC-Qwen3-8B in LIRICAL dataset. d. Hit rates of VC-Ranker and RC-Ranker across different Top K values in LIRICAL dateset.

In the PUMCH-ADM dataset, RC-Qwen3-8B demonstrated a significant predictive advantage, with Top-1 and Top-5 hit rates reaching 0.760 and 0.933, respectively, markedly outperforming the VC-strategy (**Figure 7a**). This superior performance is attributed to the high homogeneity of the PUMCH-ADM cohort, which consists of curated, high-quality clinical data from a single center. In this setting, the real-case retriever (RC-Ranker) achieved a Top-3 hit rate of 95% (**Figure 7b**), allowing real cases to provide the model with references that closely mirror the target cases. Beyond this internal comparison, the RC-Qwen3-8B result holds a more significant finding when benchmarked against external baselines. Even when employing the lightweight Qwen3-8B model, our RC-strategy surpassed the best reported outcomes from RareBench’s most capable models, including GLM3-Turbo and Gemini-Pro. This superior performance, notably achieved with a lightweight LLM, directly demonstrates the efficiency gains realized through our specialized RC-Ranker prioritization and case augmentation technique over the original RareBench implementation.

However, the trend reversed in the LIRICAL dataset, which better reflects the complexity and heterogeneity of the real world. VC-Qwen3-8B achieved a Top-1 of 0.602 and a Top-5 of 0.781, consistently surpassing the RC-strategy (**Figure 7c**). Data analysis reveals that because LIRICAL cases are derived from diverse global sources with high phenotypic heterogeneity, the effectiveness of real-case retrieval dropped significantly (Top-3 hit rate of RC-Ranker only 57%). In contrast, the virtual cases generated from public knowledge bases served as more robust and reliable “knowledge anchors” for the model, covering 96% of diseases in the LIRICAL repository (**Figure 7d**).

In conclusion, while the RC-strategy offers exceptional value in specialized medical centers with rich, homogeneous case archives, VC-strategy demonstrates superior robustness and scalability in broader clinical screening and heterogeneous diagnostic scenarios. By decoupling performance from the density of local case repositories, the virtual case-based framework provides a more universal and extensible technical path for addressing the “data sparsity” challenge in the rare disease domain. Future strategies may leverage public repositories as a universal baseline, with institution-specific private cases providing additional, non-disclosed enhancements tailored to local clinical practice.

## 4 Conclusion

This study introduces the virtual cases augmented reasoning, a novel paradigm designed to systematically bolster the diagnostic reasoning capabilities of LLMs within the rare disease domain. By synthesizing high-fidelity “Virtual Standardized Cases” directly from authoritative, expert-vetted knowledge bases such as HPO, Orphanet, and OMIM, the VC-RDAgent effectively addresses the critical limitations of existing diagnostic tools. Unlike traditional methodologies that rely on fragmented, privacy-sensitive patient records or real-time web-crawling, the VC-strategy operates entirely within a controlled, offline-capable environment. This architecture ensures a reproducible and privacy-preserving diagnostic pipeline that remains robust even when high-fidelity real-world reference cases are unavailable, thereby providing a scalable solution to the pervasive challenge of data sparsity in rare disease clinical practice.

Extensive benchmarking across diverse clinical datasets and model architectures demonstrates that the virtual cases function as a powerful, model-agnostic performance elevator. For frontier closed-source models, the framework refines the convergence of diagnostic decisions and enhances top-tier accuracy. More significantly, the application of virtual cases results in a profound harmonization of performance across varying model scales, effectively narrowing the gap between mid-size open-source models and state-of-the-art frontier systems. The finding that light-weight models, such as the 8B-parameter Qwen3, can achieve diagnostic recall parity with high-parameter counterparts, together with the observation that leading open-source models, such as GLM4.7, approach the performance envelope of commercial frontier systems, underscores the ability of virtual cases to provide essential diagnostic priors. This suggests that high-precision clinical decision support can be localized to resource-constrained environments, reducing the operational dependency on proprietary, cloud-based infrastructures while maintaining rigorous standards of clinical safety and evidence-based reasoning.

Furthermore, the comparative analysis between virtual and real-case strategies highlights the superior generalizability of the VC-strategy in managing the immense phenotypic heterogeneity of the rare disease spectrum. While real-case retrieval may offer advantages in highly homogeneous institutional datasets, virtual cases provides a more stable “knowledge anchor” in complex, real-world scenarios characterized by atypical or incomplete clinical presentations. By decoupling diagnostic efficacy from the density of local case registries, VC-LLMs achieve a level of robustness and stability that exceeds both zero-case baselines and traditional similarity-based algorithms. This stability ensures that the correct diagnosis remains within the primary clinical focus, mitigating the risk of diagnostic errors caused by idiosyncratic noise or linguistic variability in patient reports.

In conclusion, the VC-RDAgent demonstrates that the synthesis of structured virtual cases is a viable and effective strategy for enhancing the diagnostic reasoning of LLMs in the identification of rare diseases. By enabling models of various scales to navigate the clinical complexities of rare conditions, this research provides a versatile foundation for developing more accessible and reliable decision support tools. Ultimately, the framework offers a pragmatic and privacy-conscious pathway for integrating AI-driven insights into clinical workflows, addressing the persistent challenges of phenotypic heterogeneity and data scarcity within a scalable technical architecture.

## 5 AI Agent Setup

To facilitate clinical adoption by rare disease specialists and ensure strict patient data privacy, we developed a user-friendly web application for VC-RDAgent (**Figure 8**). The platform enables users to input clinical presentations and obtain diagnostic predictions through an intuitive, chat-based interface optimized for clinical workflows. The VC-RDAgent web application is accessible at https://rarellm.service.bio-it.tech/rdagent/. As reflected in the underlying implementation, the diagnostic workflow encompasses four sequential phases.

**Figure 8.**
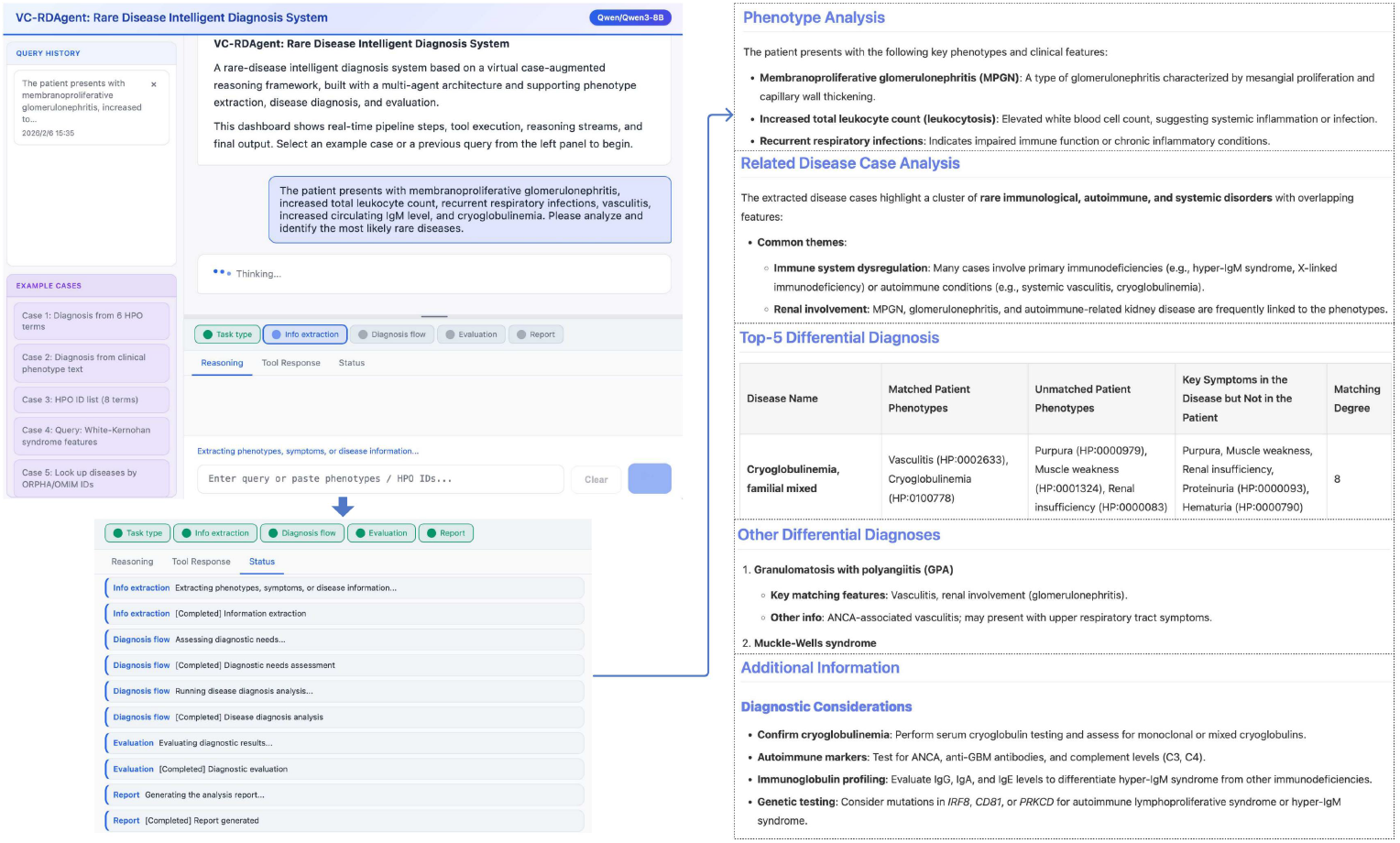
Overview of the VC-RDAgent web application workflow.

### Clinical Data Entry

Users can input free-text patient descriptions, structured symptom lists, or explicit HPO identifiers directly through the conversational interface.

### Phenotype Extraction and Standardization

Upon submission, the backend automatically parses the unstructured input and extracts relevant clinical features. The interface displays real-time status indicators (e.g., “Extracting phenotypes…”) while the system maps extracted features to standardized HPO terms using internal extraction and normalization logic, preparing the data for downstream analysis.

### Virtual Case Augmented Reasoning

Unlike systems that rely on real-time web search, the platform executes the privacy-preserving VC-LLMs workflow entirely locally. The backend orchestrates the VC-Ranker to synthesize virtual standardized cases from the internal knowledge base. In parallel, the frontend provides a traceable reasoning panel that allows clinicians to inspect intermediate reasoning steps and verify how patient phenotypes are aligned with virtual disease profiles during the chain-of-thought process.

### Evaluation and Report Generation

In the final stage, the system performs a safety-oriented consistency check on the proposed diagnostic candidates to ensure logical coherence and clinical plausibility. The system then consolidates the results into a structured diagnostic report, including sections such as phenotype analysis, related disease cases analysis, and Top-10 differential diagnosis, ensuring that the output is clinically actionable.

## Data Availability

The code and data for the VC-RDAgent project, including its implementation for rare disease diagnosis, are publicly available. This includes the source code, integrated knowledge base and related scripts. The project’s repository can be accessed at: https://github.com/cloudna-AI4LS/VC-RDAgent.

## Prompt Templates

**STEP1 DIAGNOSIS**

You are an expert in rare diseases.

A patient presents with the following detailed phenotypes:

**[structured details of patient phenotypes]**

Please follow the structured reasoning approach below to identify exactly 10 potential candidate rare diseases

**based solely on the provided phenotypes**.

**REASONING APPROACH (Strict):**

1. Use ONLY the phenotypes provided.
2. Consider clusters of phenotypes by organ/system to detect disease patterns.
3. The patient may have multiple underlying conditions; not all phenotypes need to fit one disease.
4. Treat all phenotypes as potentially relevant; do not over-prioritize any single feature.
5. Du not exclude a disease if some phenotypes are missing.
6. Consider inferred patient factors (age, sex, onset, progression) from phenotypes.
7. Prioritize rare and well-documented diseases with high phenotype specificity.

**FINAL ANSWER FORMAT (strict):**

List the 10 most likely candidate diseases in order, with no explanations, and the output must start with “FINAL ANSWER”:

FINAL ANSWER:

1. DISEASE_NAME
2. DISEASE_NAME
3. DISEASE_NAME
4. DISEASE_NAME
5. DISEASE_NAME
6. DISEASE_NAME
7. DISEASE_NAME
8. DISEASE_NAME
9. DISEASE_NAME
10. DISEASE_NAME

**STEP2 DIAGNOSIS:**

You are an expert in rare diseases.

A patient presents with the following detailed phenotypes:

**[structured details of patient phenotypes]**

The following diseases are considered highly relevant candidates (**unordered list; no ranking implied**), list with the associated phenotypes:

**[Retrieval-based virtual cases from VC-Ranker]**

**[Generation-based virtual cases from step1 output]**

Please follow the structured reasoning approach below to identify the most likely diseases for this patient.

**IMPORTANT ADDITIONAL INSTRUCTION:**

For each candidate disease, you must not only rely on the explicitly listed phenotype associations above, but also incorporate your own medical and genetic knowledge base to identify additional potential phenotype-disease associations.

This ensures that diseases are not unfairly excluded due to incomplete associations in the provided list.

If your knowledge base indicates possible links between the candidate disease and the patient’s phenotypes, you should include that in your reasoning.

**REASONING APPROACH (Strict):**

1. **List all provided phenotypes.**
2. **Cluster phenotypes by organ/system**.
3. **Identify key phenotype patterns/signatures** that indicate likely disease classes.
4. **Cross-match phenotype clusters with the provided candidate diseases.**
  - The patient may have multiple diseases; not all phenotypes must fit one disease.
  - Do not over-prioritize any single phenotype; consider all phenotypes as relevant.
  - Do not exclude a disease solely because some provided phenotypes are missing, as not all phenotypes must appear in a single disease.
  - Diseases with higher overlap of phenotype patterns should be ranked higher.
5. **Assess progression, age, and sex implications** inferred from the phenotypes.
6. **Select at least 10 candidate diseases** that best fit the phenotype distribution.

**FINAL ANSWER FORMAT (strict):**

List the 10 most likely candidate diseases in order, with no explanations, and the output must start with “FINAL ANSWER”:

FINAL ANSWER:

1. DISEASE_NAME
2. DISEASE_NAME
3. DISEASE_NAME
4. DISEASE_NAME
5. DISEASE_NAME
6. DISEASE_NAME
7. DISEASE_NAME
8. DISEASE_NAME
9. DISEASE_NAME
10. DISEASE_NAME

